# Characterization of *Escherichia coli* ExbD protein modification *in vivo*

**DOI:** 10.1101/2020.01.29.925446

**Authors:** Aruna Kumar, Kathleen Postle

## Abstract

The TonB system of *Escherichia coli* couples the protonmotive force of the cytoplasmic membrane to active transport of nutrients across the outer membrane. In the cytoplasmic membrane, this system consists of three known proteins, TonB, ExbB, and ExbD. ExbB and ExbD appear to harvest the protonmotive force and transmit it to TonB, which then makes direct physical contact with TonB-dependent active transport proteins in the outer membrane. Using two-dimensional gel electrophoresis, we found that ExbD exists as two different species with the same apparent molecular mass but with different pIs. The more basic ExbD species was consistently present, while the more acidic species arose when cells were starved for iron by the addition of iron chelators. The cause of the modification was, however, more complex than simple iron starvation. Absence of either TonB or ExbB protein also gave rise to modified ExbD under iron-replete conditions where the wild-type strain exhibited no ExbD modification. The effect of the *tonB* or *exbB* mutations were not entirely due to iron limitation since an equally iron-limited *aroB* mutation did not replicate the ExbD modification. This constitutes the first report of *in vivo* modification for any of the TonB system proteins.

---

For Gram-negative bacteria, the characteristic outer membrane (OM) poses a dilemma. It protects the cells from environmental assailants such as detergents and antibiotics, but it limits the nutrients that can be obtained by diffusion to those ~600 da or less (1). Transport of larger, scarce or very important nutrients is accomplished by a wide variety of active transport proteins in the OM that obtain the energy from the protonmotive force of the cytoplasmic membrane (CM) (2–8). Energy transduction proceeds cyclically through a complex of integral CM proteins ExbB, ExbD, and TonB, with ExbB/D harvesting the protonmotive force and transducing it to TonB protein, which then transmits the energy for active transport to a TonB-dependent transporter (TBDT) in the OM (9–11). The TBDTs consist of a 22-stranded β-barrel that is entirely occluded by an amino terminal globular domain, called the cork (12). At least one role of TonB is to energize movement of the cork to allow transport into the periplasm (13–17).

Although involved in transport of many different nutrients such as nickel, sugar, and vitamin B12, for *Escherichia coli* the main function of the TonB system is to acquire the normally biologically unavailable element iron (18, 19). In an aerobic environment at neutral pH, iron is present in virtually insoluble ferric-hydroxide complexes. Likewise, mammalian hosts keep levels of soluble iron low by the presence of iron-binding proteins such as transferrin and lactoferrin (20). To acquire iron the bacteria therefore synthesize and secrete molecules with high affinities for iron (21). These siderophores (*Gr*. iron bearer) can chelate and solubilize the iron, with the iron-siderophore complex subsequently transported across the OM, energized by the TonB system. Expression of TonB, and the unlinked operon encoding ExbB, and ExbD are regulated ~ 3-to-4-fold under aerobic conditions by the availability of iron and the Fur repressor protein (9, 22, 23). Under iron limiting conditions, the expression of TonB, and potentially therefore also ExbD, is induced as the culture grows to stationary phase due to progressive iron starvation (24). The expression of FepA, the OM transporter of the only siderophore synthesized by domesticated *E. coli* K12, is regulated over a wider range of ~ 30-fold (9). The *E. coli* K12 siderophore, enterochelin, is synthesized from chorismate as an offshoot of the pathway for aromatic amino acid biosynthesis by a non-ribosomal peptide synthetase (25). Both enterochelin (also known as enterobactin) and its biosynthetic intermediates are absent in *aroB* strains (26).

The integral cytoplasmic membrane proteins TonB, ExbB, and ExbD appear to form a complex in the cytoplasmic membrane with TonB extending across the periplasm to mechanically energize active transport through customized beta barrels in the outer membrane (11, 17, 27–29). TonB and ExbD are anchored in the cytoplasmic membrane by their N-termini, with the majority of each protein extending into the periplasmic space (Fig. 1) (30–32). With ExbB acting as a scaffold, the relationship between TonB and ExbD periplasmic domains proceeds through a four-stage cycle where the protonmotive force of the cytoplasmic membrane is required for ExbD to configure TonB for correct interaction with TonB-dependent transporters (TBDTs) (11, 17).

**Fig. 1.**
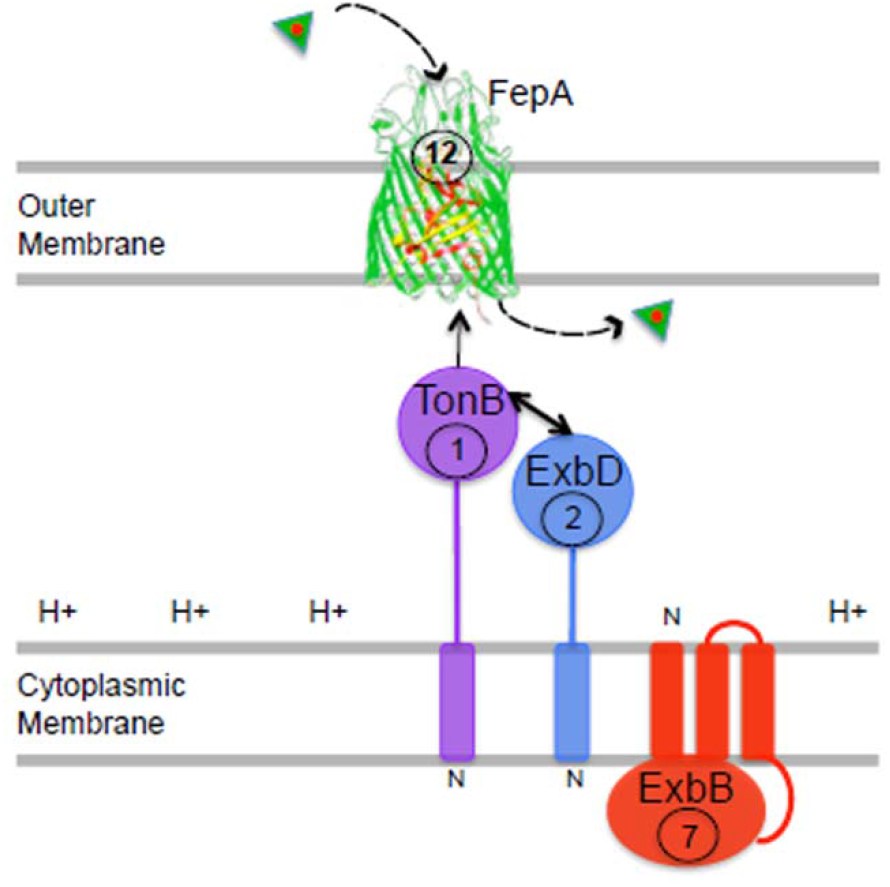
The TonB system. The siderophore enterochelin, containing an iron atom (green triangle), binds to the TonB-dependent transporter, FepA with sub-nanomolar affinity (29). Integral cytoplasmic membrane proteins TonB, ExbB and ExbD harvest the protonmotive force of the cytoplasmic membrane and somehow transduce it to FepA, causing it to release the enterochelin in the direction of the periplasmic space. From that point it is picked up by a periplasmic binding protein and carried to a cytoplasmic membrane ABC transporter for transport into the cytoplasm (49). Numbers in circles represent the per-cell ratios of each protein in *E. coli* W3110 grown under iron limiting conditions. Under those conditions, TonB is present at ~ 1300 copies per cell (9). [Adapted from (11), which also contains a model for the dynamics of the TonB system *in vivo*.]

To understand more about the role of ExbD in the energy transduction process and search for possible chemical modifications, we evaluated its behavior by 2-dimensional (2D) gel electrophoresis. Membrane protein preparations were made from cells grown as liquid cultures in minimal M9 medium supplemented with additional aromatic amino acids as described previously (13) to mid or late exponential phase and pelleted by centrifugation. The pellet was suspended in 10mM HEPES buffer at 4° C and lysed by French pressure cell 3 times at 20000 psi. All subsequent manipulations took place at 4° C until samples were prepared for gel analysis. Unlysed cells were removed by centrifugation at 1000x g. Membranes were pelleted at 40 K rpm (~140,000 x g) for 2hr. The pellet was solubilised in 1.5ml of 10mM HEPES, 0.5% Na-deoxycholic acid with shaking for 2hr. The unsolubilised material was pelleted at 29 K (~ 75,000 x g) for 1hr. Solubilised membrane proteins were analyzed on conventional 2-dimensional gels following BIO-RAD’ protocol for mini-PROTEAN II, 2D Cell. In the first dimension, ampholines in the pH range of 3-10 were used. The second-dimension gel was 13% SDS polyacrylamide (Fig 2). ExbD was detected on the 2D separation at pI range of 4.5-5.5 by immunoblot with α-ExbD polyclonal antibodies (Fig 2c), consistent with the calulated pI of ExbD at 4.7. Our initial experiment examined over-expressed ExbD from KP1038 [GM1, *exbB*::Tn*10*, *tolQ_am_*, (33)] carrying plasmid pKP390 (*exbB/D* operon under arabinose control in pBAD24) induced with 0.2% arabinose, where two different spots with the same molecular mass but different apparent pI values were detected (Table 1, Fig. 2).

**Fig. 2.**
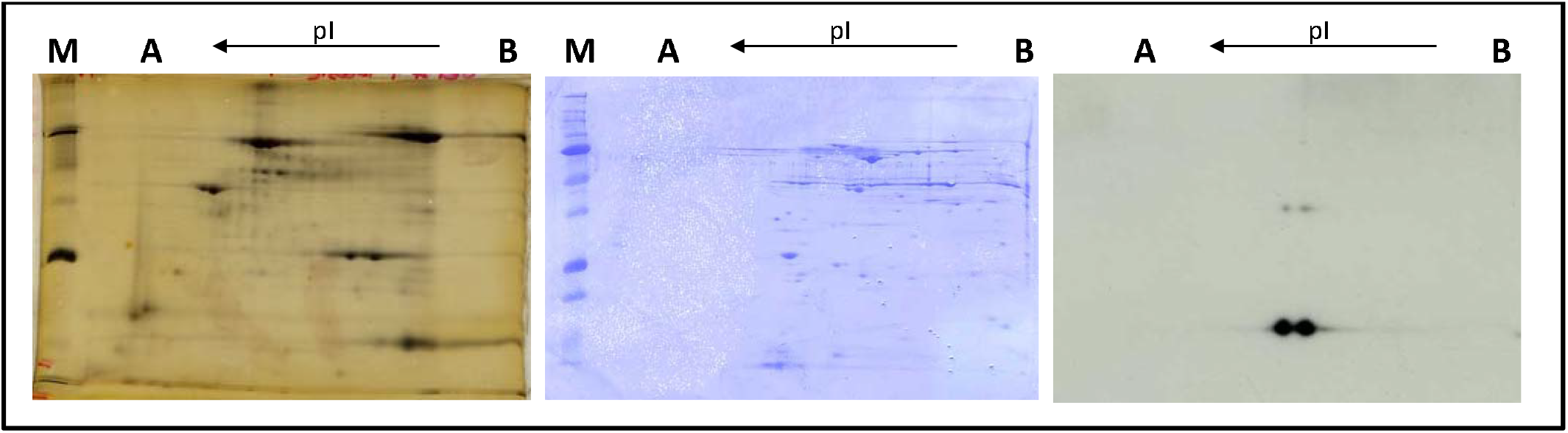
Two-dimensional (2D) gel electrophoresis analysis of ExbD protein. A) Distribution of silver-stained pI markers in descending order of molecular mass: hen egg white conalbumin (pI = 6.0, 6.3, 6.6, MW = 76,000), bovine serum albumin (pI = 5.4, 5.6, MW = 66,200), bovine muscle actin (pI = 5.0, 5.1, MW = 43,000), rabbit muscle GAPDH (pI = 8.3, 8.5, MW= 36,000), bovine carbonic anhydrase (pI = 5.9, 6.0 MW= 31,000, soybean trypsin inhibitor (pI=4.5, MW =21,500), equine myoglobin (pI = 7.0, MW = 17,500). The arrow represents the direction of isoelectric focusing from basic (B) to acidic (A). B) 2D analysis of total membrane fraction (see text) from strain GM1 grown to A_550_ = 0.85 (measured using a Spectronic 20 with a 1.5 cm pathlength) electrophoresed on 2D gels, and stained with Coomassie brilliant Blue. C) 2D analysis of ExbD protein. Immunoblot of membrane fraction separated by 2D gel electrophoresis and developed using anti-ExbD antibody (9) at 1:5000 and secondary antibody at 1:10000.

**Table 1.** Bacterial strains and plasmids used

| Strains or<br>plasmids | Properties | Source or reference |
| --- | --- | --- |
| GM1 | *ara* (*pro-lac*) *thi* F' *pro Lac* | (44) |
| W3110 | F- IN(*rrnD-rrnE*)I *rph-1* | (45) |
| KP1038 | GM1 (*exbB*::Tn10, *tolQ* ) | (33) |
| KP1092 | GM1(*fepA*::*kan*) | (46) |
| KP1274 | GM1 (*aroB zhf-50*::Tn10) | (47) |
| KP1344 | W3110 $\Delta$(*tonB*, *P14*)::*blaM*) | (35) |
| KP1422 | GM1(*fur*::Tn5) | (9) |
| KP1484 | GM1, Δ*tonB*::*kan* | This study |
| KP1487 | W3110, Δ*fepA*::*kan* | This study |
| KP1491 | W3110, $\Delta$(*tonB*, *P14*)::*kan*, Δ*fepA* | This study |
| pKP378 | *exbD* gene in pBAD24 | (48) |
| pKP390 | *exbB/D* operon in pBAD24 | (48) |

Observations on overexpressed ExbD could be subject to artifact. To determine the cause of the ExbD modification, we then completed all further studies in a strain background where ExbD was expressed in its normal chromosomal context in GM1 (*ara*, Δ*lac-pro, thi*; F’ *pro, lac*), referred to in this paper as “wild-type”. Bacterial strains were grown in M9 medium supplemented with additional aromatic amino acids as described (13) to either mid-exponential or late exponential phase under a variety of iron availabilities (Fig. 3). In the presence of added iron (FeCl_3_.6H_2_O at 1.85μM or 100μM), only one form of ExbD was evident (rows a, b; columns D, E). However, in the absence of any added iron, an apparently modified, more acidic, version of ExbD appeared when cells were grown to late exponential phase (compare column C in rows a and b) and were expected to be more iron-starved (24). The intensity of the modified more acidic ExbD spot increased as the cells were starved for iron by growth in the presence of iron chelators 2,2’-dipyridyl (DIP) at 200 μM and diethylenetriaminepentaacetic acid (DTPA) at 100 μM (Fig. 3, rows a and b, columns A and B). Thus, the degree of ExbD modification was greater when the cells were starved for iron. It was not possible to grow the strains that are intrinsically iron-limited (*aroB, fepA, tonB, tonB + fepA*, or *exbB*) in the presence of iron chelators.

**Fig. 3.**
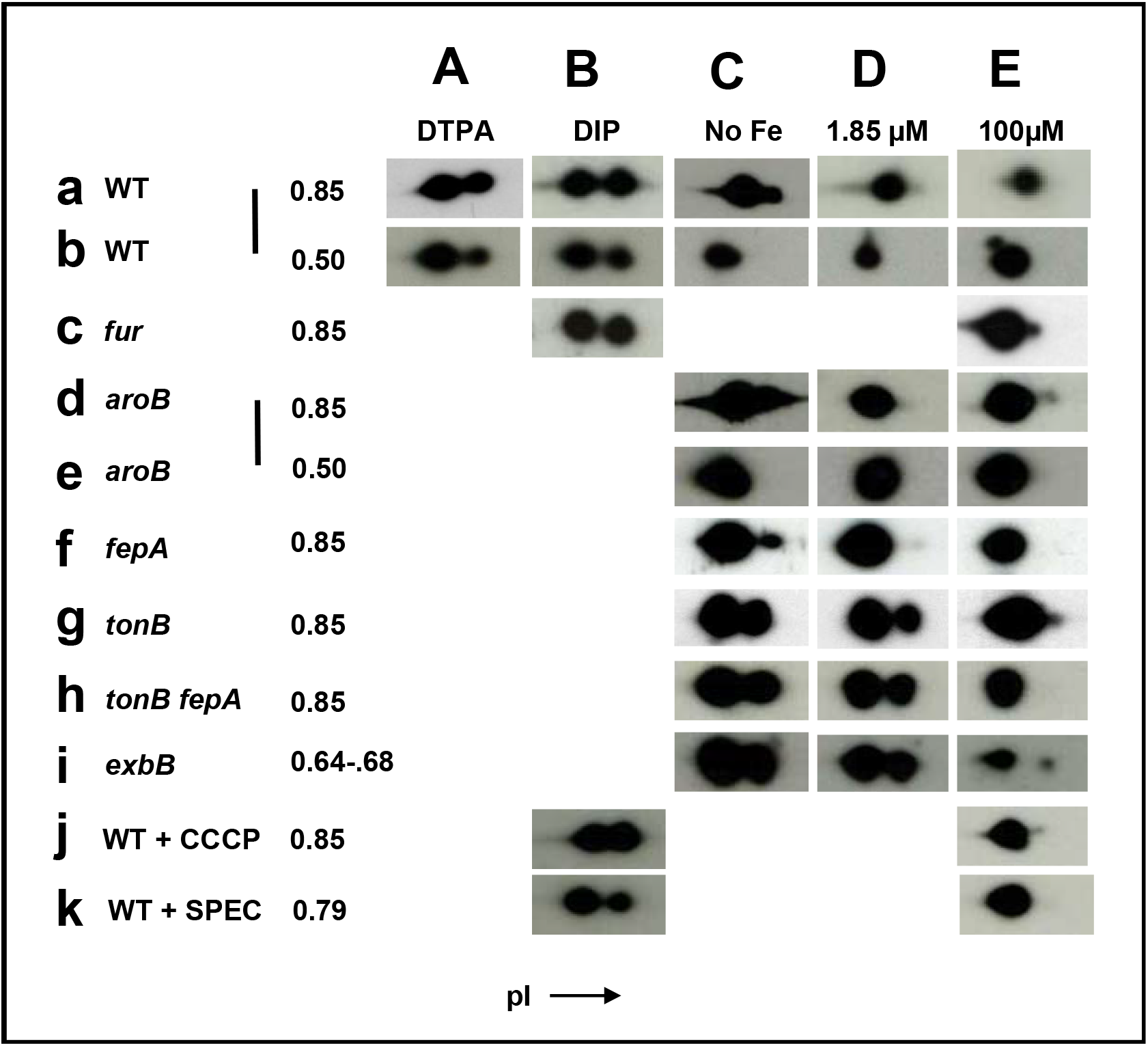
Differential modification of ExbD protein detected by 2D gel electrophoresis. Regions containing monomer ExbD-specific bands arising from the immunoblots similar to those in Fig. 1C are shown. a) and b) Strain GM1 grown either to A_550_ = 0.85 or A_550_ = 0.50; c) KP1422 (GM1 *fur*::Tn*5*) grown to A_550_ = 0.85; d) and e) KP1274 (GM1, *aroB*, *zhf-50*::Tn*10*) grown to A_550_ = 0.85 or A_550_ = 0.50; f) KP1487 (W3110, Δ*fepA*::*kan*) grown to A_550_ = 0.85; g) KP1484 (GM1, Δ(*tonB, P14*)::*kan*) grown to A_550_ = 0.85; h) KP1491 (W3110, Δ(*tonB, P14*)::*kan*, Δ*fepA*) grown to A_550_ = 0.85; i) KP1038 GM1 (*exbB*::Tn*10*, *tolQ*) carrying pKP 378 (*exbD*) was grown in the presence of 0.008% arabinose only to A_550_ = 0.64; j) GM1 grown to A_550_ = 0.85 and treated with 10μM CCCP (carbonyl cyanide 3-chlorophenylhydrazone) for 5 min prior to analysis on 2D gels; k) GM1 grown to A_550_ = 0.79 and treated with 200μM spectinomycin for 1 hr. prior to analysis on 2D gels. It is important to note that only relative levels of ExbD and modified ExbD are being compared within each panel. Comparison of levels between panels is not shown. Arrow depicts direction of isoelectric focusing from basic to acidic. A = 100μM diethylene-triamine-penta-acetic acid (DTPA); B = 200μM 2,2’ Dipyridyl (Dip); C= No supplement added; D= 1.85μM of FeCl3.6H2O; and E = 100μM of FeCl_3_.6H_2_O represent increasing degrees of iron abundance. Note that different A_550_ values were reached prior to processing for Aa (0.73), Ci (0.64), and Di (0.68), likely due to iron limitation.

Fur is a global repressor of siderophore transport genes and enterochelin biosynthetic genes, as well as *tonB, exbB* and *exbD* (23, 24, 34). The effect of a Fur mutation (strain KP1422; GM1, *fur*::Tn*5*), which can mimic the transcriptional regulatory effects of iron limitation on the TonB system (9), was determined for the ExbD modification for cells grown with high levels of iron and iron starved cells (+DIP). The finding of only one major form of ExbD in the presence of 100 μM Fe and two equally abundant forms of ExbD in the presence of DIP suggested that the effect on ExbD modification occurred downstream of its transcriptional regulation by Fur (Fig. 3, row c, columns B, E). These results further suggested that the protein which modified ExbD might not be regulated by Fur.

*In vivo* interaction of TonB with TBDT FepA is enhanced in presence of its transport ligands (17). In addition, TonB undergoes a conformational change following contact with a TBDT in the presence of enterochelin that does not occur in its absence (35). To test whether the absence of enterochelin (the sole TonB-dependent ligand synthesized by *E. coli* K12) affected ExbD modification, ExbD encoded by strain KP1274 (GM1, *aroB, zhf-50*::Tn*10*) was evaluated on immunoblots of 2D gels (Fig. 3, rows d,e; columns C, D, E). Similar results were seen for parental strain GM1 at both mid- and late-exponential phase, which ruled out the possibility that enterochelin-dependent signaling through FepA was the source of the modification. Similar results were also obtained for the strain KP1487 lacking FepA (W3110, Δ*fepA*::*kan*; Fig 3, row f).

There is evidence for signal transduction among TonB, ExbB, and ExbD in the cytoplasmic membrane. *In vivo*, ExbB and ExbD proteins are needed to generate conformational changes in the TonB C-terminus and for TonB to respond conformationally to the presence or absence of protonmotive force (10, 35, 36). The TonB N-terminal transmembrane domain is required for correct configuration of its C-terminus for subsequent interaction with ExbD (11). An intrinsically disordered region near the N-terminal transmembrane domain of ExbD regulates interaction of its C-terminus with TonB (37). To study effects of the absence of TonB, the modification of ExbD was examined in KP1484 (GM1, Δ*tonB*::*kan*) and KP1491 (W3110, Δ*tonB*::*kan*, Δ*fepA*) (Fig. 3, rows g, h). In both cases, when TonB was absent a greater degree of ExbD modification was detected in columns C and D compared to wild-type bacteria grown under the same conditions. The Δ*fepA* mutation did not detectably change the degree of ExbD modification seen in row h.

Strains lacking *tonB* or the ability to synthesize enterochelin (*aroB*) are intrinsically iron-limited. The increase in ExbD modification seen with the *tonB* mutants could therefore be simply due to differences in degree of iron limitation. To indirectly compare steady state iron levels among wild-type, *aroB*, and *tonB* strains, levels of OM protein FepA, which is iron regulated (9, 38), were assessed in the media with various degrees of iron availability (Fig. 4). Strains were grown in M9 media with different levels of iron or iron chelators and harvested for immunoblot analysis by precipitation with trichloro-acetic acid (TCA) at A_550_= 0.5. Harvesting with TCA prevents degradation by endogenous proteases that can occur even during boiling in gel sample buffer (39). While the level of FepA expression decreased in wild-type cells with the addition of increasingly greater levels of iron as seen previously (9), the levels of FepA in *aroB* or *tonB* cells remained constant and were essentially identical in all growth media. Although the *tonB* and *aroB* mutations led to similar degrees of iron limitation, only the *tonB* mutation led to increased ExbD modification (Fig. 3, in columns C and, especially D, compare rows d, f with g, h). The same result was observed for cells harvested at A_550_ = 0.85 (data not shown).

**Fig. 4.**
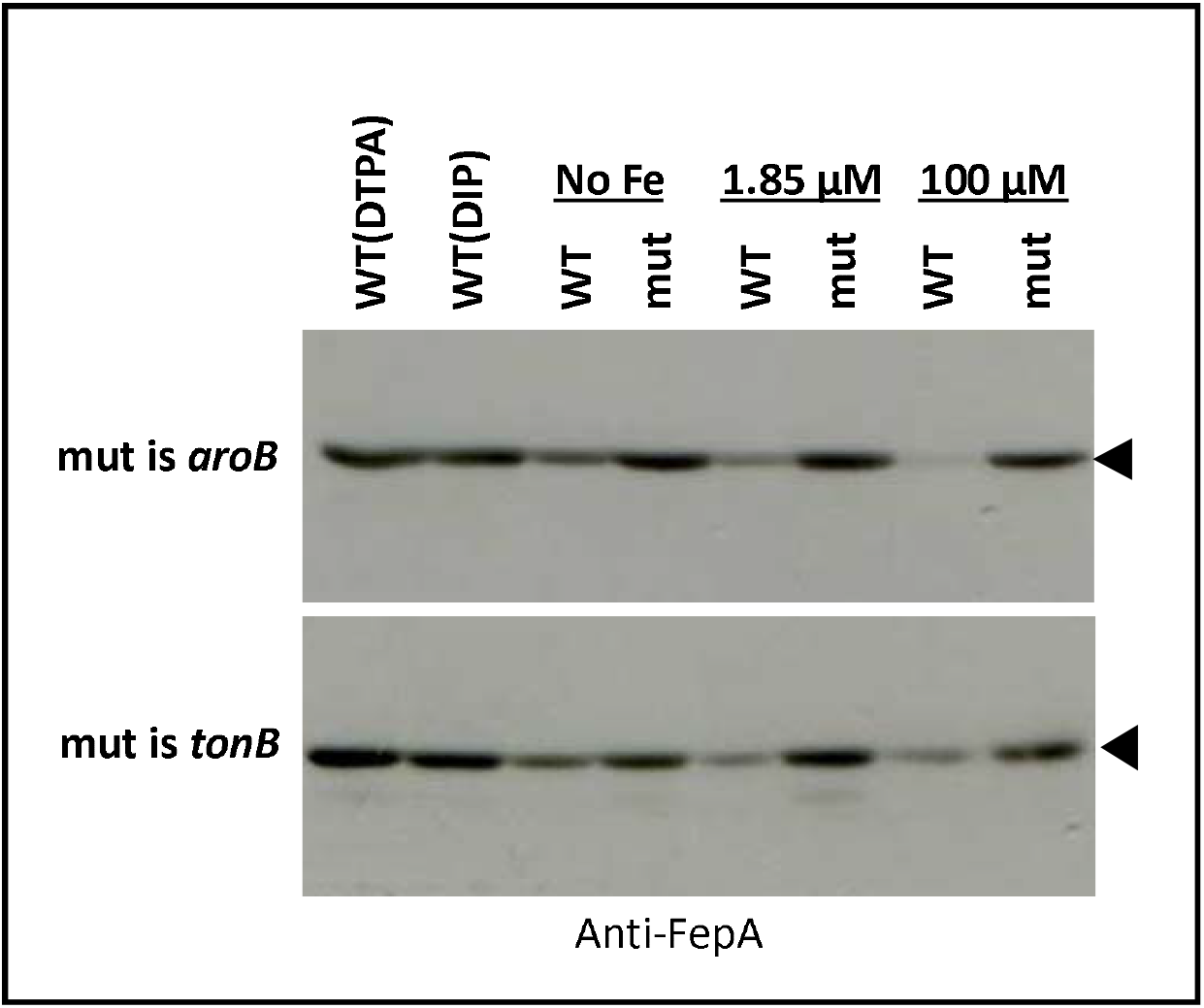
Relative FepA levels in various media are consistent with expected degrees of iron limitation. Strains were grown in minimal medium supplemented as in Fig. 3 to A_550_= 0.5, harvested by precipitation TCA. Samples were electrophoresed on 11% SDS polyacrylamide gels and analyzed by immunoblot using anti-FepA primary antibody [1:5000; (9)] and secondary HRPO conjugated sheep/Goat anti-rabbit (1: 10000). WT= GM1; mut stands for *aroB* =KP1274 or *tonB* = KP1344.

To determine the level of ExbD modification in the absence of ExbB, strain KP1038 (GM1 *exbB*::Tn*10*, *tolQ*) was used with plasmid pKP378 expressing ExbD under control of the arabinose promoter to compensate for the polarity of the Tn*10* insertion in *exbB* on *exbD* expression (23). In M9 medium with ampicillin (100 μg/ml) and glycerol as carbon source, 0.008% arabinose induced ExbD expression to levels that mimicked those of a chromosomally encoded *exbD* gene. In the absence of iron or in 1.85 μM Fe, strains grew slowly and were harvested at A_550_ = 0.64 or 0.68 respectively. Similar to the Δ*tonB* strains, the absence of ExbB resulted in significantly higher levels of modified ExbD in both the absence of any iron and in 1.85 μM iron (Fig. 3, row i, columns C, D). Wild-type strains grown to A_550_ = 0.5 or 0.8 (Fig. 3, rows a, b columns C, D) did not exhibit this increased ExbD modification, thus ruling out growth-phase as a relevant variable. Thus, it appeared that the increased modification could be due to changes that occurred within the TonB/ExbB/ExbD complex during the energy transduction cycle.

We determined if agents known to disrupt the TonB system changed the degree of ExbD modification. The TonB complex responds conformationally to the presence or absence of protonmotive force (10, 11, 35). To determine if collapse of the protonmotive force played a role in ExbD modification, the protonophore carbonyl cyanide 3-chlorophenylhydrazone (CCCP) was added at 50 μM to GM1 cells at A_550_ = 0.85 for 5 min at 37 °C with shaking and the cells were processed for the 2D gel analysis of ExbD. 50 μM CCCP completely and immediately stops bacterial growth and TonB system activity (40). There was no change compared to wild-type cells in iron-limited media or iron-replete media (Fig. 3, compare row j, columns B, E with row a, columns B, E). Therefore, the ExbD modification appeared to occur even in the absence of TonB system function, as it also did in the absence of TonB and ExbB.

In the absence of continued protein synthesis, all the proteins in the TonB system have half-lives of ~ 120 min, while TonB activity as measured by various indirect assays decays with a half-life of ~ 20 min (11, 33, 41). This disparity suggests the presence of additional, as yet unknown, proteins in the TonB system since protein synthesis inhibitors have no effect on protonmotive force (Skare and Postle, unpublished observation). To determine if the effect of protein synthesis inhibition on ExbD modification, GM1 cells were grown in M9 medium to A_550_ = 0.7. Spectinomycin was added at 200 μg/ml and the cells were incubated for 1 hour at 37 °C (final A_550_ = 0.79). There was no significant change in the extent of ExbD modification compared to wild-type cells in iron-limited media or iron-replete media (Fig. 3, row k, columns B, E; compare to row b, columns A, B, E).

It seemed likely that the modification of ExbD was occurring through its periplasmic domain, since there are only ~ 20 residues predicted to occupy the cytoplasm and those could be sequestered by ExbB [Fig. 1; (11)]. Phosphorylation of periplasmic proteins has been documented (42), however, in that case it did not appear to be of functional significance (43). Neither ExbD nor its modified form could be detected by immunoblots with anti-phosphoserine, anti-phosphotyrosine, or anti-phosphothreonine antibodies (data not shown), suggesting that the modification was either not phosphorylation or that it did not occur through those residues.

To the best of our knowledge, this is a first report of a chemical modification of any component of the TonB system. We have not determined the role it plays, however we thought it was sufficiently important to future research to publish the data so that others can take note of them.

## Acknowledgements

We thank Charles Bulathsinghala for construction of KP1484 and Surendran Devanathan for construction of KP1487 and KP1491. We thank Michael Konkel for instruction in the 2D gel electrophoresis technique. We thank Anne Ollis and Ray Larsen for critical reading of the manuscript. We gratefully acknowledge the support of the National Institute of General Medical Sciences through grant GM42146 to K.P.

